# Newly developed silicone airway stent (GINA stent): *Mechanical characteristics and performance evaluation in pigs*

**DOI:** 10.1101/2020.10.24.343533

**Authors:** Hwa Sik Jung, Ganghee Chae, Jin Hyoung Kim, Soohyun Bae, Chui Yong Park, Soyeoun Lim, Soon Eun Park, Moon Sik Jung, Ju Ik Park, Young Jae Lee, Sung Kwon Kang, Don Han Kim, Yongjik Lee, Taehoon Lee

## Abstract

**OBJECTIVES:** Central airway obstruction (CAO) is caused by various malignant and benign processes. Surgery is a preferred option for CAO, but if not possible, bronchoscopic treatment could be performed. Recently, bronchoscopic treatments have been improved. Particularly in airway stents, new attempts are being made to overcome the existing shortcomings of stents (migration, mucostasis, and granulation tissue formation). We recently developed a new silicone airway stent (GINA stent). The GINA stent has anti-migration design, dynamic structure enabling reduction of stent cross-sectional area, and radiopaqueness. We sought to evaluate mechanical characteristics and performance of our novel GINA stent in a pig tracheal stenosis model.

**METHODS:** All tests were performed by comparing GINA stent [outer diameter (OD, mm) 14, length (L, mm) 55] with Dumon stent (OD14L50). Mechanical tests were done using digital force gage to determine the anti-migration force, expansion force, and flexibility. Short-term (3 weeks) performance was evaluated after stent implantations [GINA (n = 4) vs. Dumon (n = 3)] in a pig model of tracheal stenosis.

**RESULTS:** Mechanical properties outcomes for GINA vs. Dumon: anti-migration force [18.4 vs. 12.8 Newton (N)]; expansion force (11.9 vs. 14.5 N); flexibility (3.1 vs. 4.5 N). Short-term (3 weeks) GINA vs. Dumon performances: mucus retention (0/4 vs. 0/3); granulation tissue formation (0/4 vs. 0/3); migration (1/4 vs. 2/3).

**CONCLUSIONS:** GINA stent demonstrated better mechanical properties than Dumon stent with a stent performance not inferior to Dumon stent.

## INTRODUCTION

The central airway obstruction (CAO) is broadly defined as a blockage of the trachea, either main stem bronchus and/or the bronchus intermedius, and it might be caused by various malignant and benign processes.[1] Treatment of CAO, surgery (resection and anastomosis) could provide the best opportunity for definitive management. However, because of anatomical limitations, or advanced and/or metastatic disease, or poor medical conditions, surgery treatment of CAO is limited to specific benign processes such as tracheal stenosis after infection, intubation, or tracheostomy. After all, most CAOs could only be solved by bronchoscopic interventions, which requires mostly rigid bronchoscopy, and includes balloon dilatation, mass excision (debulking, debridement), tumor ablation (cryotherapy/argon plasma coagulation/electrocautery/laser), and stent insertion.[2] While it is desirable to resolve CAO without a stent, in reality, a stent is the primary technique utilized in CAO treatment, owing to the cartilage-damaged benign stricture or externally compressive malignant stricture that occupies most of the CAO.[3, 4]

There are two main types of stents, each with different characteristics. The metal stent has a strong expansion (radial) force, which has the advantage of a low migration rate. However, it has a risk of granulation tissue overgrowth and tracheobronchial perforation. Silicone stent, on the other hand, has a small expansion force, thereby, minimizing the risk of granulation tissue overgrowth and tracheobronchial perforation, but migration occurs readily. Mucus clogging occurs well in both types of stents.[1] As such, airway stent is effective for immediate relief of CAO but causes complications such as migration, mucostasis, and granulation tissue formation frequently. To overcome these shortcomings, stent technologies are evolving persistently, with some invented for specific purposes. Examples are drug-eluting stent (for granulation inhibition), a stent with a new design (for migration/granulation inhibition), a stent with an inner surface hydrophilic coating (for mucostasis inhibition), radiopaque silicone stent, customized three-dimensional (3-D) printed stent, and bio-absorbable stent.[2, 4–6]

We recently developed the new silicone airway stent (named “GINA stent”) based on anti-migration design, with a flexible, dynamic structure to enable a reduction of the stent cross-sectional area to enhance bronchial expiratory flow, as well as radiopaque capability for readily accessible radioimaging. The purpose of the present study was to introduce GINA stent by learning more about it mechanical and performance quality in comparison with an already established Dumon stent via anti-migration force, expansion force, and flexibility determinations, along with an evaluation of short-term (3 weeks) performance in a pig tracheal stenosis model.

## MATERIALS AND METHODS

### The GINA stent: a newly developed silicone airway stent

GINA stent was made of biocompatible medical-grade silicone through a process of injection molding. GINA stent had three characteristics. First, <u>anti-migration design</u>: outer ring (right-angled triangle shape) of GINA stent was designed to fit into the cartilaginous trachea; the plane of GINA stent that comes into contact with the membranous trachea was characterized by a raised three-line arrangement. Second, we considered <u>a flexible, dynamic structure to facilitate the reduction of the stent cross-sectional area</u>: GINA stent maintains physiological airway contraction via the flat portion of the stent (like the membranous portion of tracheobronchial), which makes it more flexible and helps to remove airway secretions through an enhanced expiratory flow. Third, <u>radiopaqueness</u>: barium sulfate was added to the silicone to make the stent radiopaque to be easily identifiable in radiological imaging studies (Figure 1).

**Figure 1.**
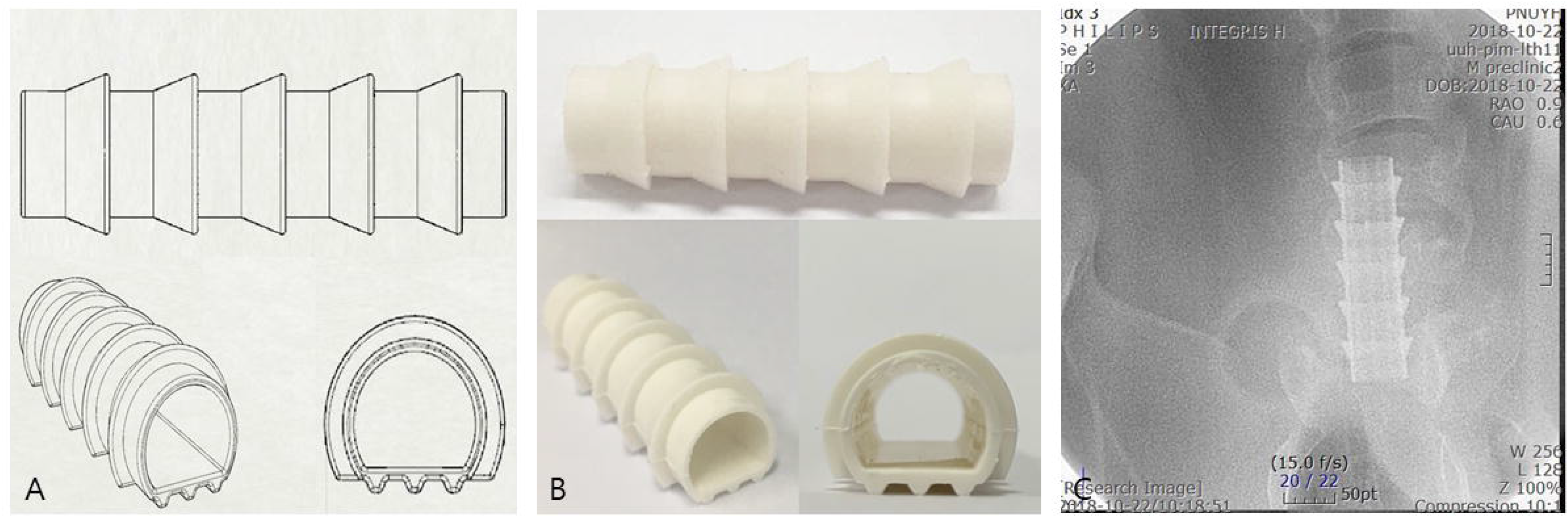
GINA stent, the silicone stent recently developed for this study. The GINA stent has an anti-migration design, dynamic structure enabling a reduction of the stent cross-sectional area, and radiopaqueness capability. (A) Design of GINA stent. (B) Actual GINA stent [outer diameter of 14 mm (OD14) / ring diameter of 18 mm (RD18) and length of 55 mm (L55)]. (C) Radiopaqueness of GINA stent.

All measurements performed in this study were made by comparing the GINA stent with Dumon stent (Novatech, France). For the mechanical tests and animal study, GINA stent with an outer diameter of 14 mm (OD14), ring diameter of 18 mm (RD18), length of 55 mm (L55), and a wall thickness of 1 mm and Dumon stent with an outer diameter of 14 mm (OD14), surrounding studs of 2 mm (which makes the longest diameter of Dumon stent 18 mm), length of 50 mm (L50), and a wall thickness of 1.5 mm were selected.

### Mechanical tests of GINA and Dumon stents

Three mechanical properties (anti-migration force, expansion force, and flexibility) were tested by Research and Development Department of S&G Biotech (Gyeonggi-do, Korea), a well-known stent maker in Korea (www.sngbio.co.kr), using the following methods. The measurement of forces was performed by the digital force gage (LR5K Plus, Lloyd Instruments, Hampshire, England).

The anti-migration force was measured by the following method. The Teflon jig with an inner diameter of 16 mm was fixed to the tensile strength tester. After inserting the stent into the Teflon jig, the push gage was used to move the stent 5 cm to the opposite side of the Teflon jig. At that time, the value of the anti-migration force was measured (Figure 2A). At least three measurements were made for either stent. The highest value was selected as an anti-migration force of each stent. For GINA stent, both forward and backward directions were performed.

**Figure 2.**
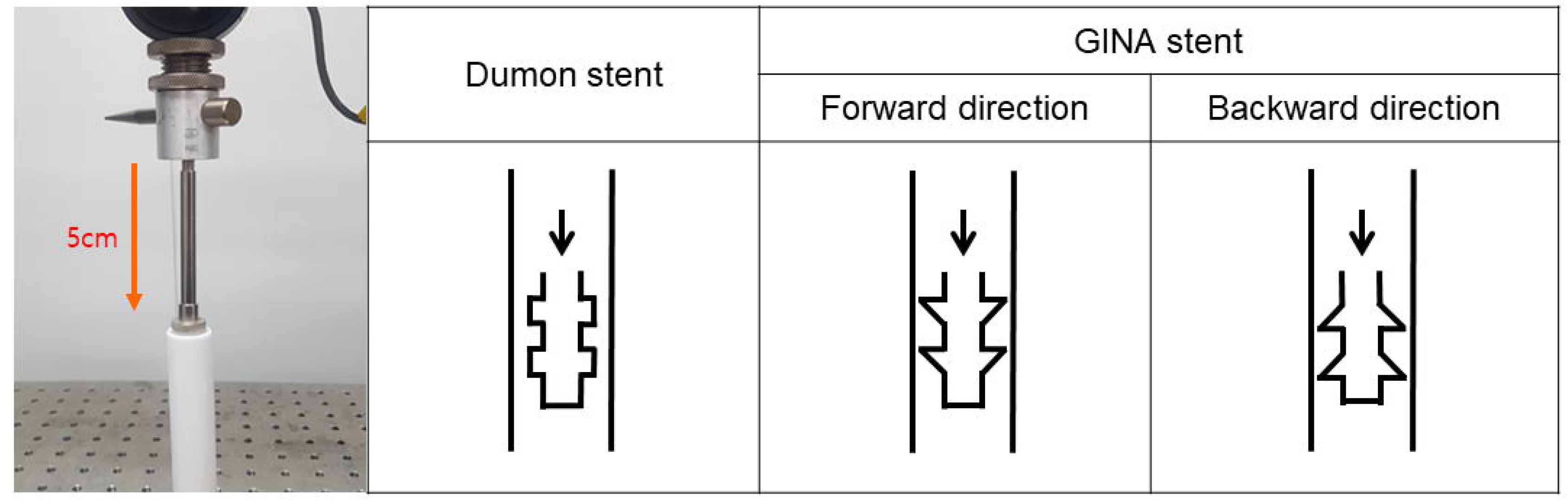
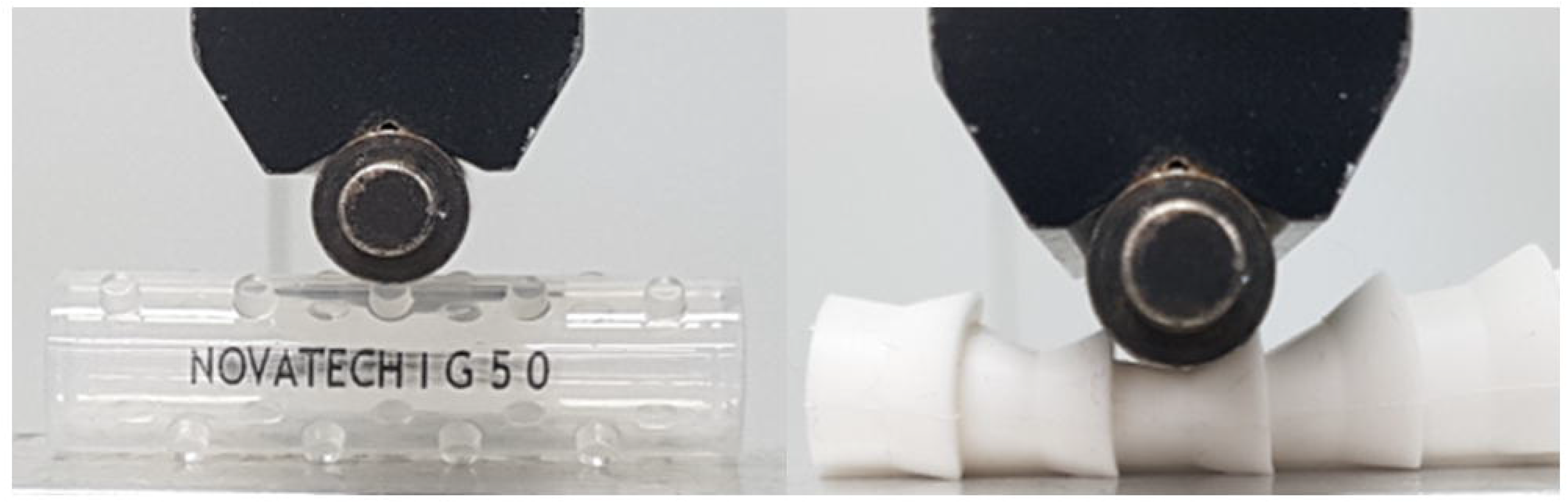
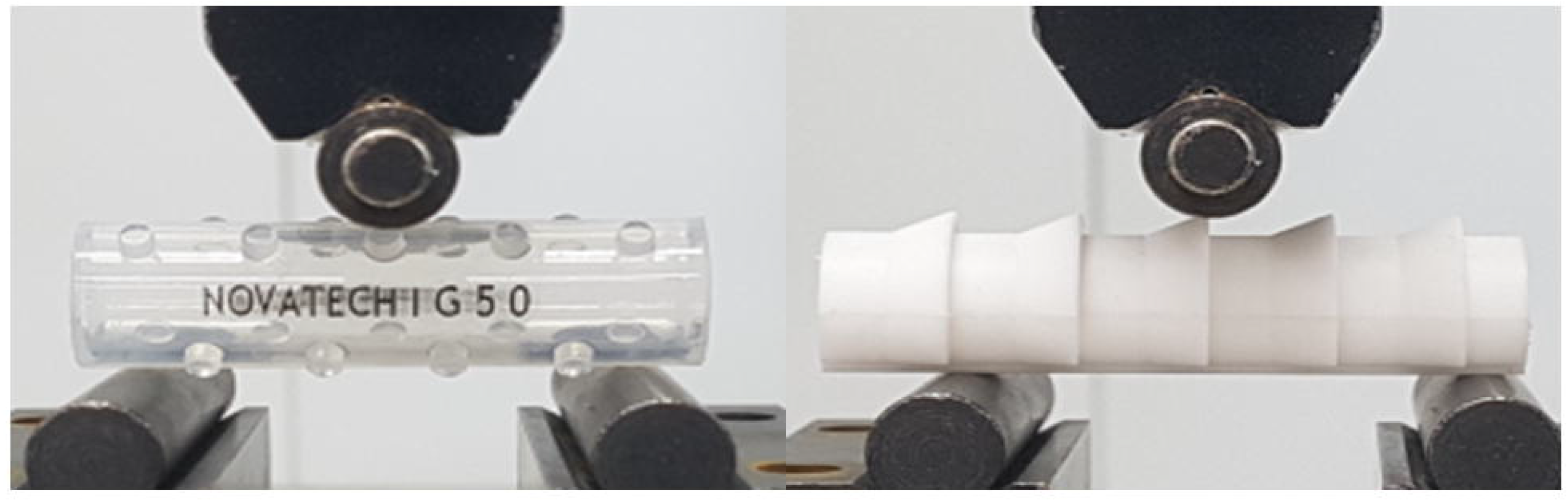
The mechanical test methods. Three mechanical properties (anti-migration force, expansion force, and flexibility) of GINA and Dumon stents were measured by digital force gage. (A) Anti-migration force was measured by pushing (5 cm) a stent through in a Teflon jig. In the case of GINA stent, both forward and backward directions were performed. (B) Expansion force was measured by pushing a stent on a flat surface until the stent diameter was reduced by 50%. (C) Flexibility was measured by pushing a stent on separated jigs until the stent diameter was reduced by 50%.

The expansion force was measured as follows. After placing the stent on a flat floor, the measuring load cell was attached to the stent surface. The force was measured until the diameter of the stent was reduced by 50% (Figure 2B). The measurements were made in various directions, repeating at least three tests and recordings in each direction. The highest value among different directions was selected as an expansion force of each stent.

The flexibility measurements were performed, as follows. After separating the measuring jigs at intervals of 4 cm, the stent was put on the jigs. The measuring load cell was attached to the stent surface. The initial force was set to zero. The measuring load cell pushed the stent slowly and proceeded to half the diameter of the stent. The force was measured, according to the advance of the measuring load cell (Figure 2C). Measurements were made in various directions, repeating at least three tests and recordings in each direction. The lowest value among different directions was selected as the flexibility of each stent.

### Animal performance study in GINA and Dumon stents

Short term (3 weeks) performance assessment of GINA stent, in comparison with Dumon stent, was carried out in a pig model of tracheal stenosis. For this experiment, 12 weeks-old seven female farm pigs (body weight, 40–45 kg) were selected because of the similarity in tracheal size (16–20 mm) pigs share with humans.

Tracheal stenosis was made by the method of cuff overpressure intubation (COI) and tracheal cautery (TC), which we developed recently.[7] Briefly, 200mmHg of COI via a silicone tracheal tube (internal diameter (ID) 9.0 mm/outer diameter (OD) 12 mm) was applied to the pig’s trachea for 1 hour. Immediately after COI, cautery (40 watt by coagulation suction tube (10390 BN, Karl Storz, Germany)) was performed on the tracheal mucosa where the cuff was located via a rigid bronchoscope (size 8.5, 10318 BP, Karl Storz, Germany) (Figure 3A). After 7 days of COI and TC, stent insertion was performed following a confirmation of induction of tracheal stenosis, revealed by a reduction of ≥50% of the measured airway cross-sectional area (Figure 3B).

**Figure 3.**
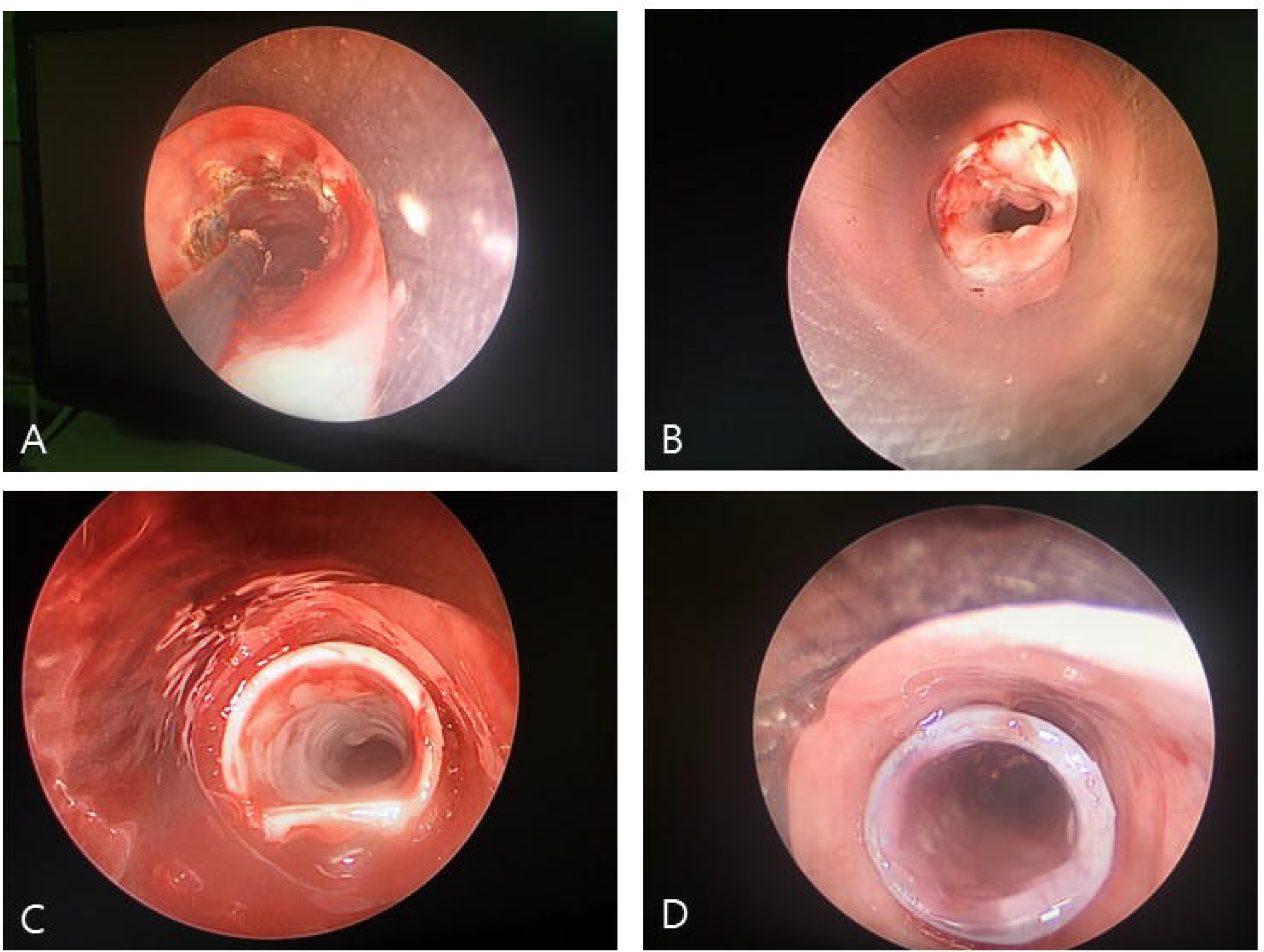
The animal study of GINA stent comparing with Dumon stent. After inducing tracheal stenosis in a 12-week-old pig, a stent (GINA or Dumon) was inserted and observed for 3 weeks. (A) Stenosis induction by tracheal cautery (TC) immediately after cuff overpressure intubation (COI). (B) Induced tracheal stenosis (a reduction of at least 50% of the measured airway cross-sectional area). (C and D) Inserted GINA and Dumon stents in the stenotic trachea of pigs.

Among 7 pigs, GINA stent was applied to 4 pigs, and Dumon stent was applied to 3 pigs (Figure 3C and 3D). TONN/NOVATECH stent applicator (Novatech, France) was used for stent positioning via a rigid bronchoscope (size 14, 10318 GL, Karl Storz, Germany). Stent performance was examined on a weekly basis by flexible bronchoscopy (MAF-TM, Olympus, Tokyo, Japan) for 3 weeks after stent insertion. The stent performance was checked for migration, granulation tissue overgrowth at both ends, and mucostasis.

For the general anesthesia, intramuscular alfaxan (5 mg/kg) and inhalational 3% isoflurane were used for induction and maintenance, respectively. This study protocol was approved by the Institutional Animal Care and Use Committee at Pusan National University Hospital (Approval Number: PNUYH-2017-043).

## RESULTS

### Mechanical characteristics of GINA stent

The anti-migration forces are illustrated in Figure 4A. Compared to the Dumon stent, the GINA stent showed a stronger anti-migration force in both directions [Dumon stent: 12.8 Newton (N) vs. GINA stent: 15.2 N (in the forward direction) and 18.4 N (in backward direction)]. The stronger the anti-migration force, the lower the probability of stent migration, thereby minimizing the inherent complications caused by this implantation technique.

**Figure 4.**
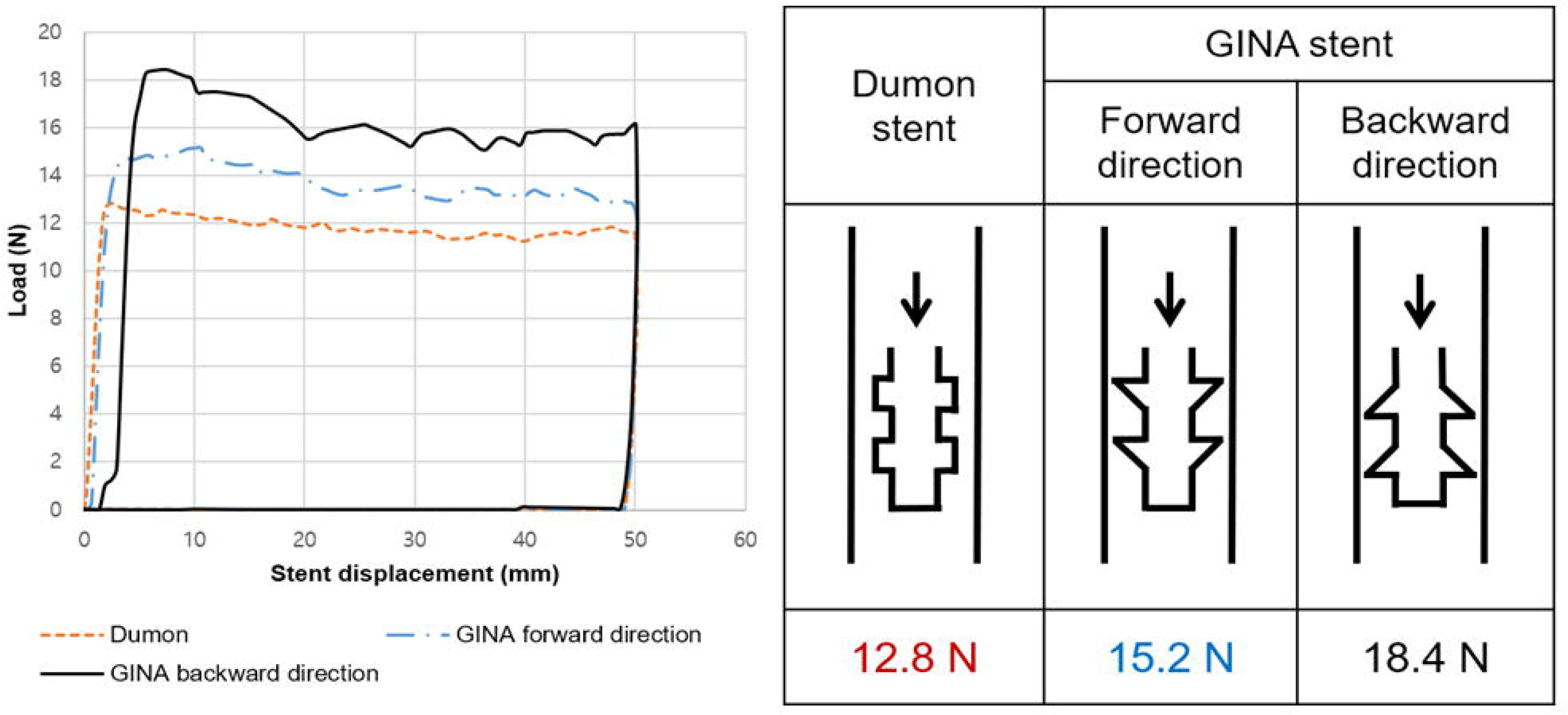
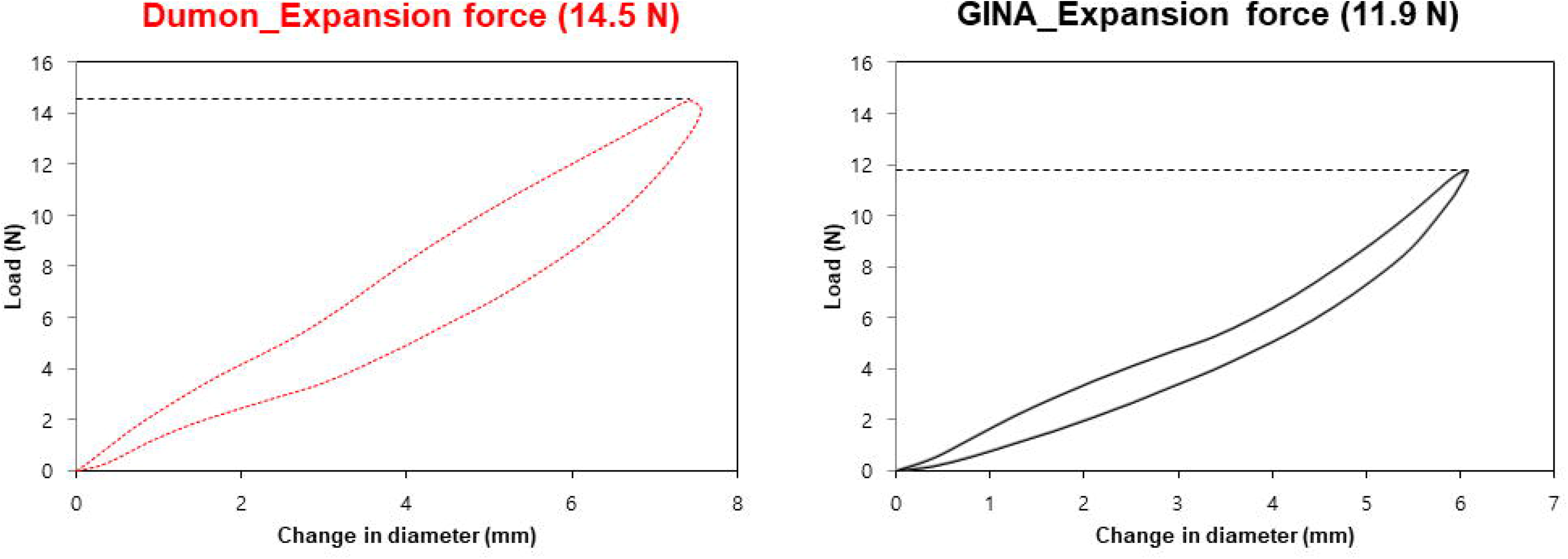
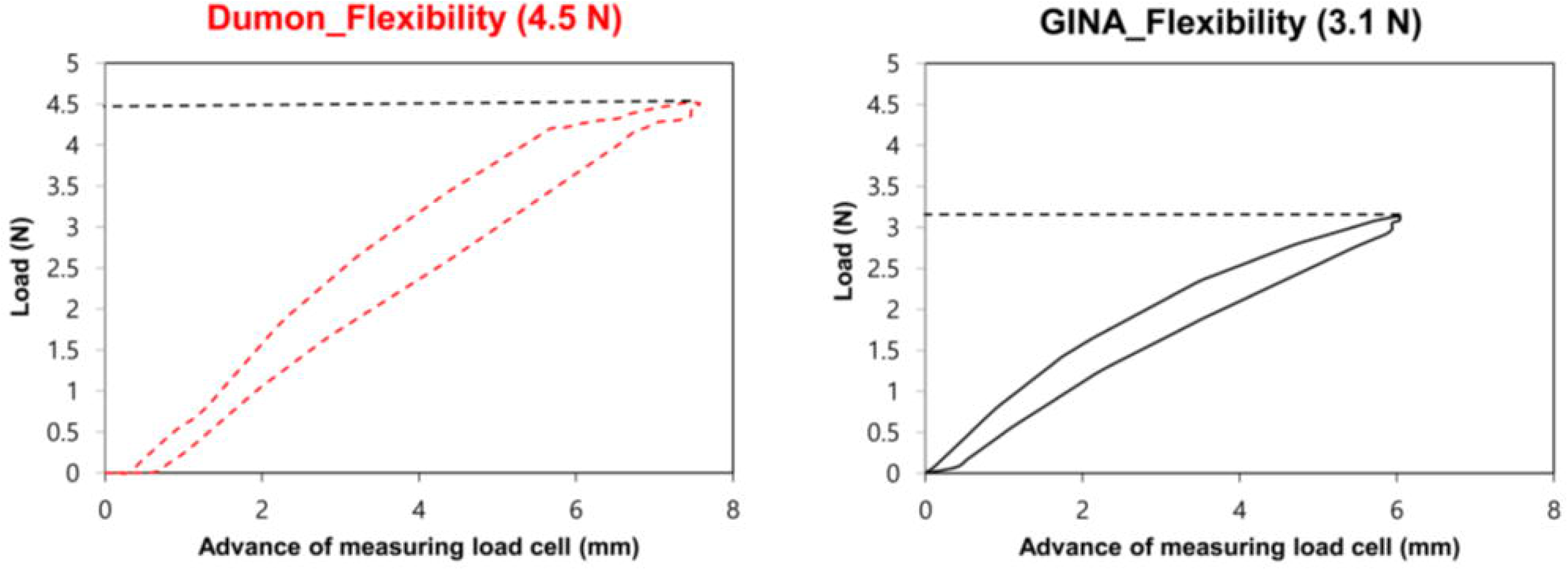
The results of the mechanical tests. (A) Compared with Dumon stent, the GINA stent showed stronger anti-migration force in both directions. (B and C) GINA stent showed a lower expansion force and higher flexibility.

The GINA stent showed a lower expansion force than Dumon stent (11.9 N vs. 14.5 N) (Figure 4B). Additionally, in terms of stent flexibility, GINA stent required a reduced force to flex the stent than Dumon stent (3.1 N vs. 4.5 N) (Figure 4C). When the expansion force is low, and the flexibility is high, the risk of granulation tissue formation and airway perforation could be lowered. Through the results of our mechanical tests, it is apparent that GINA stent has superior mechanical properties, compared to Dumon stent.

### Performance of GINA stent in a pig model

Short-term (3 weeks) performance test results of GINA stent, in comparison with Dumon stent in a pig model of tracheal stenosis, are summarized in Table 1. Stent migration was observed in one of the four pigs with GINA stent insertion, which was identified on the 14th day after stent implantation. Whereas, among three pigs with Dumon stent, two encountered stent migration on the 7th and the 14th day after stent insertion. Mucus retention and granulation tissue formation did not occur in both types of stents for the 3 weeks.

**Table 1.**
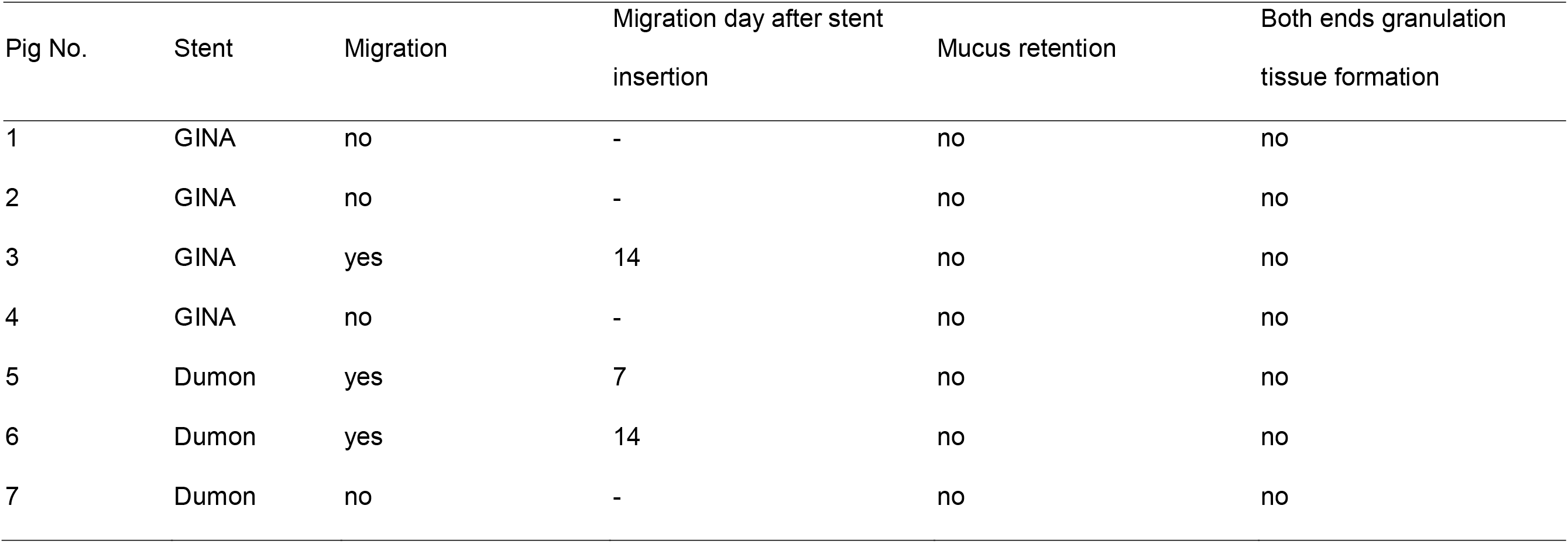
Short-term (3weeks) performance of GINA stent comparing with Dumon stent in a pig model of tracheal stenosis.

| Pig No. | Stent | Migration | Migration day after stent<br>insertion | Mucus retention | Both ends granulation<br>tissue formation |
| --- | --- | --- | --- | --- | --- |
| 1 | GINA | no | - | no | no |
| 2 | GINA | no | - | no | no |
| 3 | GINA | yes | 14 | no | no |
| 4 | GINA | no | - | no | no |
| 5 | Dumon | yes | 7 | no | no |
| 6 | Dumon | yes | 14 | no | no |
| 7 | Dumon | no | - | no | no |

## DISCUSSION

GINA stent, a radiopaque silicone airway stent we recently developed, was designed to minimize migration, mucostasis and formation of granulation tissue. The mechanical tests confirmed that GINA stent has a lower possibility of migration and granulation tissue formation than Dumon stent. In addition, it was suggested through animal experiments that the performance of GINA stent was not inferior to that of Dumon stent.

Conditions of ideal airway stent are cost-effectiveness, ease of insertion and removal, without migration or granulation tissue formation, not high but enough expansion force against airway stenosis, adequate flexibility to persevere airway physiology, and without impairment of mucociliary clearance.[8] However, no stent is able to satisfy all of these conditions, and if one characteristic is superior, another feature tends to be degraded; for example, if the expansion force of stent is low and the likelihood of granulation tissue formation is reduced, the airway fixation of the stent is less so the risk of stent migration increases. For this reason, the metal stent has less migration rate but the formation of granulation tissue is high, and the silicone stent has a high migration rate but less granulation tissue formation rate.[1] In terms of inhibition of mucostasis in the stent, it is desirable to maintain the mucociliary clearance, and it is necessary to be flexible so that the stent inner diameter is sufficiently reduced during exhalation. The uncovered metal stent could best preserve the mucociliary clearance. However, it is difficult to remove the stent due to epithelialization, and there is a problem of tissue ingrowth within the stent. For this reason, an uncovered metal stent is not recommended for benign airway stenosis;[9] it is limitedly suitable for palliation of malignant airway stenosis, but tumor ingrowth should an awareness.[10] After all, in order to solve mucostasis, it is most reasonable to improve the flexibility of the stent (enabling reduction of the stent cross-sectional area) and removal of airway secretions through the enhanced expiratory flow.

In the present study, GINA stent showed lower expansion force and higher flexibility than Dumon stent. Although the formation of granulation tissue in the animal model experiments did not differ, it is likely due to the short observation period. Granulation tissue formation is a common complication of silicone airway stents, although less in metal stents. [1, 3, 11] Excessive expansion force and low flexibility were predisposing factors for granulation tissue formation.[11–14] Small expansion force and high flexibility of GINA stent meant that lesser force was needed for the stent to expand and bend, with resultant lesser pressure on the airway, leading to a diminished mucosal inflammation and granulation tissue formation.

Despite the low expansion force, the GINA stent showed a higher anti-migration force compared to Dumon stent, which was also confirmed through the performance experiments. This superiority comes from the creative surface design of GINA stent, consisting of a right-angled triangle shape outer ring for cartilaginous trachea and a raised three-line arrangement for membranous trachea. Migration is a common complication of airway stent, especially, silicone stent.[15, 16] Various attempts have been made to inhibit stent migration. The Montgomery T tube was created with a side arm passing through a tracheostomy, which gives the stent a fixation to the trachea.[17] Recently, an external fixation method was introduced, which is very useful and has solved the cosmetic problem of the Montgomery T tube.[18] However, these methods are only applicable to upper tracheal stenosis. In the case of lower tracheal or bronchial stenosis, a bifurcation stent might aid in migration prevent, but its insertion was a challenge due to the size of the stent being excessively large, compared with the segment of stenosis.[19, 20] After all, in order to prevent migration of the stent, it is necessary to improve the friction (i.e., anti-migration force) by appropriate sizing [diameter larger than the stenosis, but slightly smaller (80-90%) than the airway diameter around the stenosis][3, 21] as well as improving the stent surface design (like studs, spikes, or protruding arcs).[6, 18, 22–24]

Another important feature of the GINA stent is the flexible dynamic structure enabling the reduction of stent cross-sectional area. GINA stent has a flat part, just like the membranous portion of a real trachea, which makes the stent more contractible and aid in removing airway secretions through an enhanced expiratory flow. Improvement of mucostasis by flattening a part of the stent has already been demonstrated by Freitag and Kim.[25–28] In our performance experiments, mucostasis was not appreciably different between the two stent types, which might be ascribed to the short observational period.

Lastly, GINA stent is radiopaque, which makes stent-tracking easier. The radiolucency of silicone stent (such as Dumon stent) has been considered a major drawback. Efforts have been made to improve this. Recently a radiopaque product has been released for the Dumon stent.

Despite the success found with our GINA stent development, our study does not preclude limitations. First, in the performance experiments, the number of animals used was small, and the observation period was short. Second, the mechanical tests used in our study were based on the advice of the stent company (S&G Biotech, Gyeonggi-do, Korea) and existing studies;[29, 30] thus, they were not validated. Therefore, these limitations should be considered when interpreting our results.

In conclusion, through strategic design, we have developed a new stent that has significantly reduced migration, despite lowering the expansion force and increasing flexibility to reduce the likelihood of granulation tissue. In the future, clinical trials are needed to demonstrate the efficacy and safety of GINA stent in humans.

## Table of abbreviations

3-D: Three-dimensional
CAO: Central airway obstruction
COI: Cuff overpressure intubation
ID: Internal diameter
L: Length
N: Newton
OD: OD Outer diameter
RD: Ring diameter
TC: Tracheal cautery

